# Small RNA profiling in *Mycobacterium* insights into stress adapt ability

**DOI:** 10.1101/2021.07.09.451870

**Authors:** Yingyu Chen, Wenjun Zhai, Kailun Zhang, Tingting Zhu, Li Su, Luiz Bermudez, Huanchun Chen, Aizhen Guo

## Abstract

Mycobacteria would encounter a number of environment changes during infection, and respond to it using different mechanisms. sRNA is a posttranscriptionally regulatory system for the function of genes and has been investigated in many other bacteria. Here, we used *Mycobacterium tuberculosis* and *Mycobacterium bovis* BCG infection models and sequenced the whole bacterial RNAs before and after host cells infection. Comparison of differential expressed sRNAs, by using GO and KEGG, and target predication, was carried out. Six pathogenically relevant stresses, drug resistance test, growth rate and morphology were used for screening and identify sRNAs. From these data, we identified a subset of sRNAs that are differentially expressed in multiple infection groups and stress conditions. We found that many of them were associated with lipid metabolism. Among them, ncBCG427, was significantly down-regulated when BCG entered into macrophages, and was associated with increase of biofilm formation and changed in drug susceptibility. Then, reduction of virulence possibility depends on regulating lipid metabolism.

## Introduction

*Mycobacterium tuberculosis* (*M. tb*) is the leading cause of human tuberculosis (TB), being in the top of 10 most important causes of death globally (WHO 2020). *Mycobacterium tuberculosis* complex, especially *M. tb* and *Mycobacterium bovis* (*M.bovis*), the leading cause of animal tuberculosis (Sun, et al. 2019), are responsible for major economic losses and represent a great threat to public health. Although zoonotic tuberculosis is an old disease, afflicting both humans and animals for thousands of years, it still remains a major health emergency.

Small bacterial RNA (sRNA) are posttranscriptional regulators that have been identified playing very important roles in translation and/or mRNA stability during different bacterial infections (Gong and Klumpp 2017; Jorgensen, et al. 2020). The number of investigations about the roles of sRNA in infection are increasing, although mycobacterial sRNA associated with macrophage infection, and particularly *M. tb*, has not been much addressed.

Pathogenicity of *M. tb* can be affected by many aspects, like regulating the virulence factors of bacterial (Bychenko, et al. 2021), influence the expression of host genes (Madhavan, et al. 2021), and control the cross talk between pathogen and host cells (Sun, et al. 2020). *M. tb* cell wall contains a large amount of complex lipids, which present as major effector molecules that interact with the host, modulating its metabolism and stimulating the immune response (Gago, et al. 2018). In *M. tb*, around 250 genes are involved in lipid metabolism; They not only regulate the replication and persistence of the *M. tb* inside the host cells but also influence the in cellular signaling, membrane microdomain organization and dynamics, and membrane trafficking (Rameshwaram, et al. 2018). Therefore, lipid metabolism represents an important component in the life cycle of *mycobacterium*.

The ability to adapt to diverse stresses in host environments is essential for a successful *mycobacterium* infection. When entering cells, the pathogen will certainly encounter large environmental challenges (Bansal, et al. 2017). Since in slow-growing mycobacteria, RNA and protein synthesis are slow, as compared with rapid growing microbes, one assumes that a mechanism must exists to facilitate the adaptation to rapidly changing host conditions. Given the relative paucity of information on the identity and function of sRNAs during mycobacteria infection, we used RNA-seq to comprehensively identify sRNAs differentially expressed by both *M. tb* and *M.bovis* BCG, in the intracellular and extracellular environments, as well as under six relevant stress conditions. Of interest, from a subset of sRNAs identified as differentially expressed, many of them were engaged in lipid metabolism. One of these, ncBCG427, not investigated upon bacterial taken up into THP-1 cells, deduced during exposure to iron starvation, lower pH stress, and membrane stress. We identified the ability of enhances the biofilm growth, impact drug resistance and reduce bacterium virulence, and then, speculate about that possibility depends on regulate the lipid metabolism.

## Materials and Methods

### Strains and growth conditions

*M. tb* strain 1458 (GenBank accession no: NZ_CP013475.1), originally obtained from a cow, was isolated and stocked in this laboratory. *M.bovis* Bacillus Calmette-Guérin (Tokyo Strain, ATCC 35737) was kindly provided by Junyan Liu from Wuhan University. *Mycobacterium smegmatis* MC^2^ 155 strain (*Mycobacterium smegmatis* MC^2^ 155, NC_008596.1) was donated by Luiz Bermudez, Oregon State University.

All *M. tb* related experiments in this research were carried out strictly in accordance with the biosafety-related operating procedures in the Animal Biosafety Level 3 Laboratory (ABSL-3) of the State Key Laboratory of Agricultural Microbiology in Huazhong Agricultural University. *M. tb* strain 1458 and *M.bovis* BCG were cultured to mid-log phase in 20 ml of Middlebrook 7H9 medium, supplemented with 10% oleic acid, albumin, dextrose and catalase medium and 0.05% Tween 80, and incubated at 37°C for 2 weeks.

### Cell culture

THP-1 cells were kindly provided by Chuanyou Li from Beijing Tuberculosis and Thoracic Tumor Research Institute, and were cultured in RPMI-1640 medium supplemented with 10% heat-inactivated fetal bovine serum. THP-1 cells were seeded in tissue culture plates and treated with phorbol-ester (PMA 5 μM) for 24 h to induce maturation. Cells were seeded at 90% confluence.

A549 human typeⅡ alveolar epithelial cells were kindly provided by Luiz Bermudez, Oregon State University, and were maintained in DEME medium supplemented with 10% heat-inactivated fetal bovine serum. A total of 10^5^ cells were added to each well of a 24-well tissue culture plate.

### Bacterial infection

PMA-differentiated THP-1 cells were infected with *M. tb* strain 1458 or *M.bovis* BCG at a MOI of 10 for 12 h. Supernatants contained bacterial were collected as extracellular bacterial control.

Remained cells were gently washed twice with PBS. Following washing, THP-1 cell cultures were maintained in RPMI-1640 medium and then scrapped from the flask, and collected 6 h and 24 h PI by centrifugation at 200×g for 10 min at 4°C.

### RNA extraction and sequencing

Extracellular bacterial total RNA was extracted using TRIzol reagent (Invitrogen, USA) followed the manufacturer’s instructions. For intracellular bacterial sample, bacteria ware separated from the eukaryotic cell, at the infection time points, washed twice with RPMI-1640 medium and added GTC lysis solution. After rapid lysing, the preparation was centrifuged at 5,000×g for 20 min, followed by 10,000×g for 20 seconds after GTC resuspension. Then, Tween-80 was added in 2 min and then RNA free water. Then the total RNA of intracellular bacterial was extracted using FastRNA Blue Kit (MP Biomedicals, Shanghai, China), according to the manufacture’s instruction. The concentration of total RNA was measured by Ultraviolet Spectrophotometer at 260nm/280nm (Thermo, USA). cDNA was synthesized and sent to sequence after quality control test.

### Data analysis and annotation

FASTX toolkit was used to get clean reads; then the clean reads were aligned to *M. tb* 1458 (NZ_CP013475) and *M.bovis* BCG (NC_012207) genomes respectively by Bowtie2 (Langmead and Salzberg 2012) with no more than 1 nt mismatch. Aligned reads with more than one genome locations were discarded. Uniquely localized reads that (1) 40 to 500 bp peaks; (2) median height of a single peak is no less than 20 nt; and (3) the maximum height of a single peak should not be less than 60 nt were screened.

Rfam database (http://rfam.xfam.org/) and BRSD database (http://kwanlab.bio.cuhk.edu.hk/BSRD/) were used to annotate all the sRNAs. DAVID (https://david.ncifcrf.gov/home.jsp) was used for GO enrichment analysis.

KOBAS version 3.0 (http://kobas.cbi.pku.edu.cn/) was used for KEGG pathway enrichment analysis IntaRNA(http://rna.informatik.uni-freiburg.de/IntaRNA/Input.jsp), CopraRNA (http://rna.informatik.uni-freiburg.de/CopraRNA/Input.jsp) and TargetRNA2 (http://cs.wellesley.edu/~btjaden/TargetRNA2/index.html) tools were used to predict sRNA target genes. Candidate genes predicted by all three tools were selected for further verification.

### Stress culture model

Iron starvation, carbon hunger, acidification pressure, oxidation pressure, membrane pressure model and granuloma-like models were used in this study as previously described (Gerrick, et al. 2018; Chen, et al. 2019). Briefly, iron-starvation medium was additionally treated with Chelex-100 (10g in 1000ml culture medium) to remove residual iron. Carbon hunger medium was PBS solution; pH 4.5 was for acidification pressure; 1mM tert-Butyl hydroperoxide (tBHP) or 0.05% SDS was added in 7H9 medium respectively for oxidative stress and membrane stress model. Granuloma-like models was 7H9 medium with 0.3M dextrose and pH6.0, and cultured with no oxygen.

After centrifuged 3,800 r/min for 10min, bacteria were washed in PBS and re-suspended in 1:1 stress- culture (V/V), then incubated in 5% CO_2_ for 24h. Bacterial was collected for further use.

### Real-time PCR (RT-PCR) analysis

RNA was extracted and reverse-transcribed to cDNA using HiScript II Q RT SuperMix for qPCR(Vazyme, China) or All-in-One™ miRNA First-Strand cDNA Synthesis Kit (Genecopoeia,US). *sigA* gene or *MysA* was used as an internal reference. The primers for amplification of selected genes are listed in table S1. The RT-PCR reactions were performed in 10 μl reaction mixtures including 5 μl of 2×SYBR Green Master Mix (Vazyme, China), 10 μM of forward and reverse primers, 0.2 μl Dye 2 and 1.5μl cDNA template (1:10 diluted), dd H_2_O to final 10μl. Samples and standards were run in triplicate in an ABI 7500 thermocycler (Applied Biosystems, USA) and analyzed using model 7500 SDS software v 1.3.1). PCR was carried out at 94°C for 10 min, then 94°C for 15s and 60°C for 1 min, for a total of 40 cycles.

### Overexpression of bacterium constructs

The rrnB promoter (between −200 and −8) was amplified from *M. smegmatis* and used to replace Hsp60 promoter in pMV261, by digesting the DNA with XbaⅠ-HindⅢ to generate pMV261-PrrnB. Seamless cloning technique was used to combine rrnB promoter and Term Transcriptional terminator.

All primers used in this study was listed in Table S2 using following systems: 98°C for 5 min, then 98°C for 30s, anneal 30 s min and 72°C for 2 min, for a total of 30 cycle, 72°C for 10 min. The PCR-generated fragments were cloned into a pMV261-PrrnB vector encoding kanamycin resistance. The resulting plasmids were propagated into DH5α *Escherichia coli* bacteria and then electroporated into *Mycobacterium smegmatis* MC^2^ 155. Transformants were selected on Middlebrook 7H11 agar plates containing 50 μg/ml of kanamycin and screened by PCR using primers originally used to PCR. The obtained fragment was then sequenced.

### Growth curve, morphology and drug resistance of recombinant strains

Recombinant strains were cultured in 7H9 medium with 50μg/ml kanamycin in 37° C, and taken every 12h for 7 days to draw the growth curve.

The morphology of single colony was observed and recorded under stereomicroscope. The area of each single colony was calculated by three-point rounding method. Five to 10 single colonies were recorded in each plate. Blade azure color rendering method was used to calculate MIC of all recombinant strains.

### Biofilm-forming ability testing

The recombinant strains were cultured to log phase (OD_600nm_ = 0.6 ~ 0.8), the bacterium were collected after centrifuge and resuspended and dispersed in fresh 7H9, adjusted to OD_600nm_=1. Then the bacterium were diluted with 1:100 in fresh Sauton medium with kanamycin, and inoculated into 12 well plate, sealed with sterile membrane, and incubated at 37°C for 5-7 days, the biofilm formation thickness of each strain was recorded. Three repeat groups were set up for each strain.

After remove the liquid and bacteria outside the biofilm, completely dry the plate add 1% crystal violet dye for 10 min; totally dry the plate after completely washing, add 95% alcohol for 10 min, after fully dissolving, read at OD_570nm_ after 3 times of dilution, and set three repeat holes. The differences of biofilm formation thickness and absorbance were compared.

### Invasion assay

A549 cells were infected with live recombinant bacterium at MOI of 10 for 1h at 37°C, gentamycin at 100 μg/ml to kill the extracellular bacterium for 2h. 0.1% Triton X-100 was used for lysate the cells after adequate washing. Then cell lysates were plated onto 7H11 agar plates and allowed to grow for 3-5 days for CFU counting.

### Statistical analysis

In vitro experiments were repeated at least three times. The Student’s t-test was used and when analyzing more than 2 groups, ANOVA analysis was used for two groups. In addition T-Test was used to calculate the results of the mice study. A value of p < 0.05 was considered to be significant. Graph Pad Prism version 7.0 software was used for statistical analysis.

## Results

### Comparison of the ncRNA (non-coding RNA) from extracellular and intracellular bacterial

The analysis of the differential ncRNAs in difference bacterial were compared. Totally, there were 490 ncRNAs in BCG, compared with 349 in *M. tb1458*. 247 (50.4%) ncRNAs were expressed both intracellular and extracellular in BCG, 76 (15.5%) were expressed only intracellular and 171 (34.9%) only expressed extracellular; 133 (38.1%) ncRNAs were expressed both intracellular and extracellular in *M. tb1458*, 158 (45.3%) were expressed only in cells compared with 58 (16.6%) were express only extracellular.

Totally, 190 antisense ncRNAs and 53 intergenic ncRNAs were successfully mapping in both two BCG groups; compared to 91 antisense ncRNAs and 42 intergenic ncRNAs were successfully mapping in two *M. tb* 1458 groups. 323 ncRNAs including 254 antisense and 69 intergenic ncRNAs were unique expressed in intracellular BCG, and 417 ncRNAs including 323 antisense ncRNA and 94 intergenic ncRNAs were unique in extracellular BCG. In intracellular *M. tb* 1458 strain, there were totally 297 ncRNAs, including 223 antisense ncRNA and 74 intergenic ncRNA. In extracellular *M. tb* 1458 strain, there were totally 193 ncRNAs, including 126 antisense ncRNA and 67 intergenic ncRNA. After compare the length distribution of difference ncRNA group, we found that for 0-200nt length ncRNA, in BCG group, extracellular had much more ncRNAs than intracellular group, but in *M. tb* 1458 strain, extracellular had much lower numbers (Fig 1A). Antisense and intergenic ncRNA length was then determined. Data showed that ncRNA less than 200nt, both antisense and intergenic ncRNA were more than intracellular ncRNAs in BCG. *M. tb* 1458 group took the reverse results, and had lower intergenic ncRNA in extracellular bacterial. Compared with BCG ncRNA than focus on less than 150nt, *M. tb* 1458 ncRNAs were mainly less than 100nt (Fig 1 BC).

**Fig.1.**
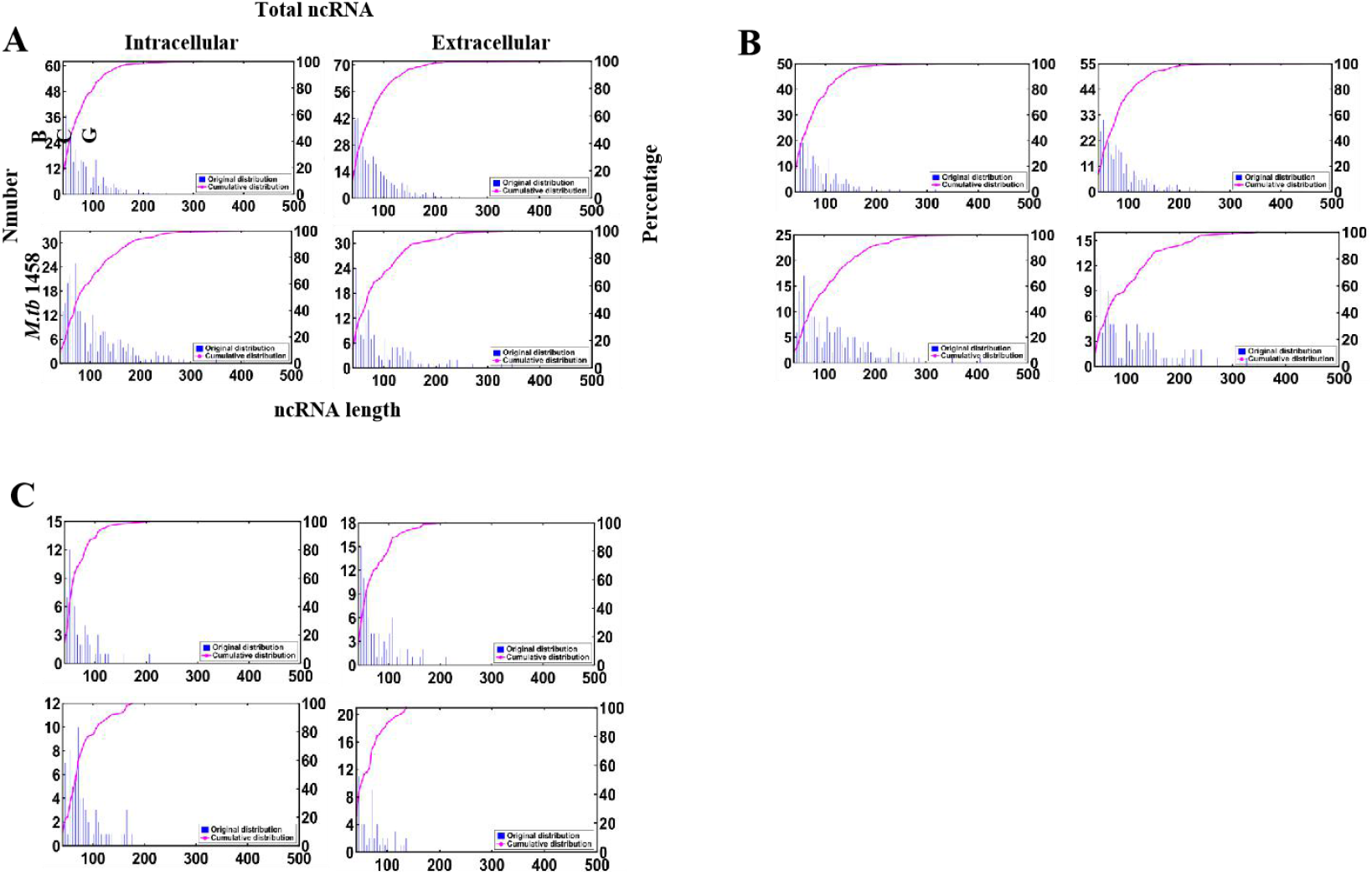
Distribution of ncRNA length. (A) Total ncRNA; (B) Antisense ncRNA; (C) Intergenic ncRNA.

### Differential expression ncRNAs in different bacterial groups

Fold change no less than 1.5, or existed only in either extracellular bacterial or in intracellular was regarded as differential expression ncRNAs in extracellular and intracellular bacterial. RPKM was used to compare ncRNAs in different groups. Compared with intracellular bacterial, in extracellular BCG, 192 ncRNAs was up regulated and 123 were down regulated; in extracellular *M. tb* 1458, 82 ncRNAs were up regulated and 177 down (Fig.2A).

**Fig.2.**
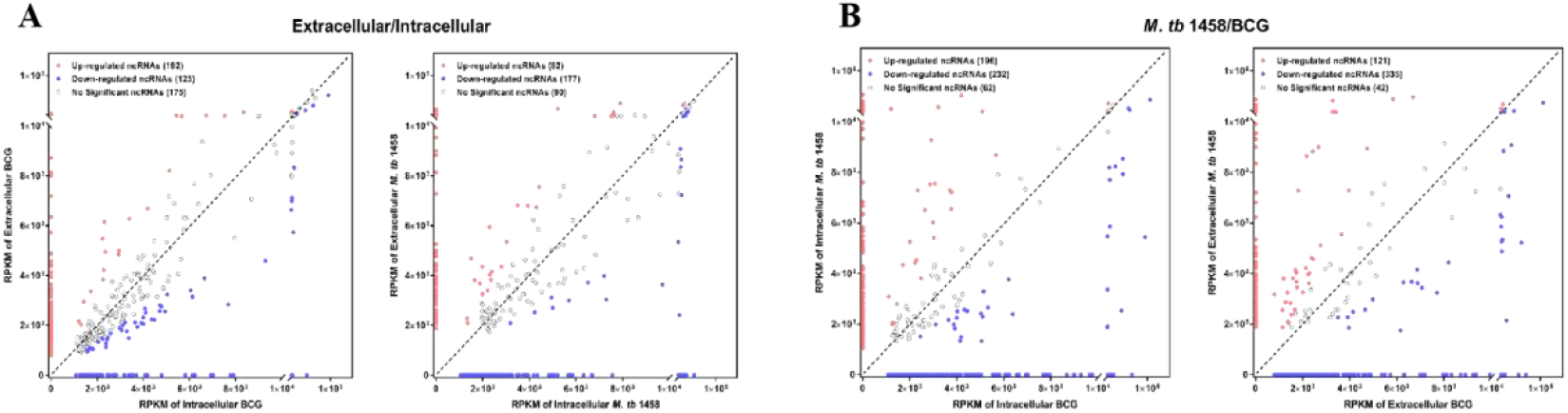
RPKM distribution of ncRNAs. (A)M. tb 1458/BCG RPKM distribution of ncRNAs between intracellular and extracellular bacteria (|FC|≥1.5). (B) RPKM distribution of ncRNAs between BCG and M. tb 1458 (|FC|≥1.5).

Fold changes no less than 1.5, or existed only in either BCG or in *M. tb* 1458, were regarded as differential expression ncRNAs between BCG and *M. tb* 1458. RPKM was used to compare ncRNAs in different groups. Compared with intracellular BCG, in intracellular *M. tb* 1458, 196 ncRNAs were up regulated and 232 were down regulated; Compared with extracellular BCG, in extracellular *M. tb* 1458, 121 ncRNAs were up regulated and 335 down (Fig.2B).

Among those differential ncRNAs, 68 was conserved in BCG and 43 in *M. tb* 1458. There were more up regulated ncRNAs in extracellular BCG than *M. tb* 1458 (Fig 3).

**Fig. 3.**
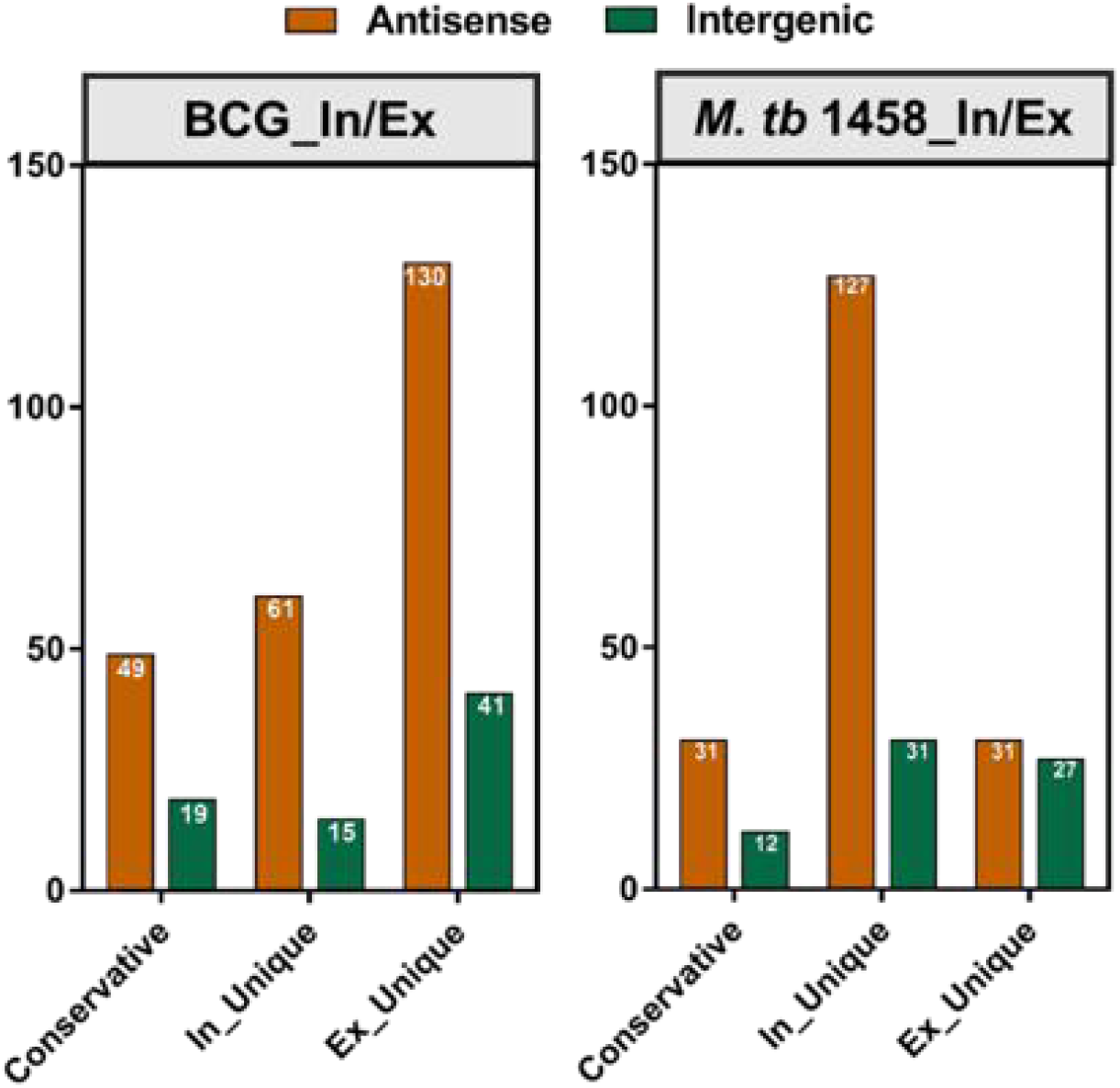
M. tb 1458/BCG counts distribution of differential ncRNAs in different types between intracellular and extracellular bacteria (|FC|≥1.5).

Compared with BCG and *M. tb* 1458, 73 were expressed in all groups, 18 ncRNAs were expressed in extracellular BCG and intracellular *M. tb* 1458; 5 only expressed in intracellular BCG and extracellular *M. tb* 1458 (Fig. 4).

**Fig. 4.**
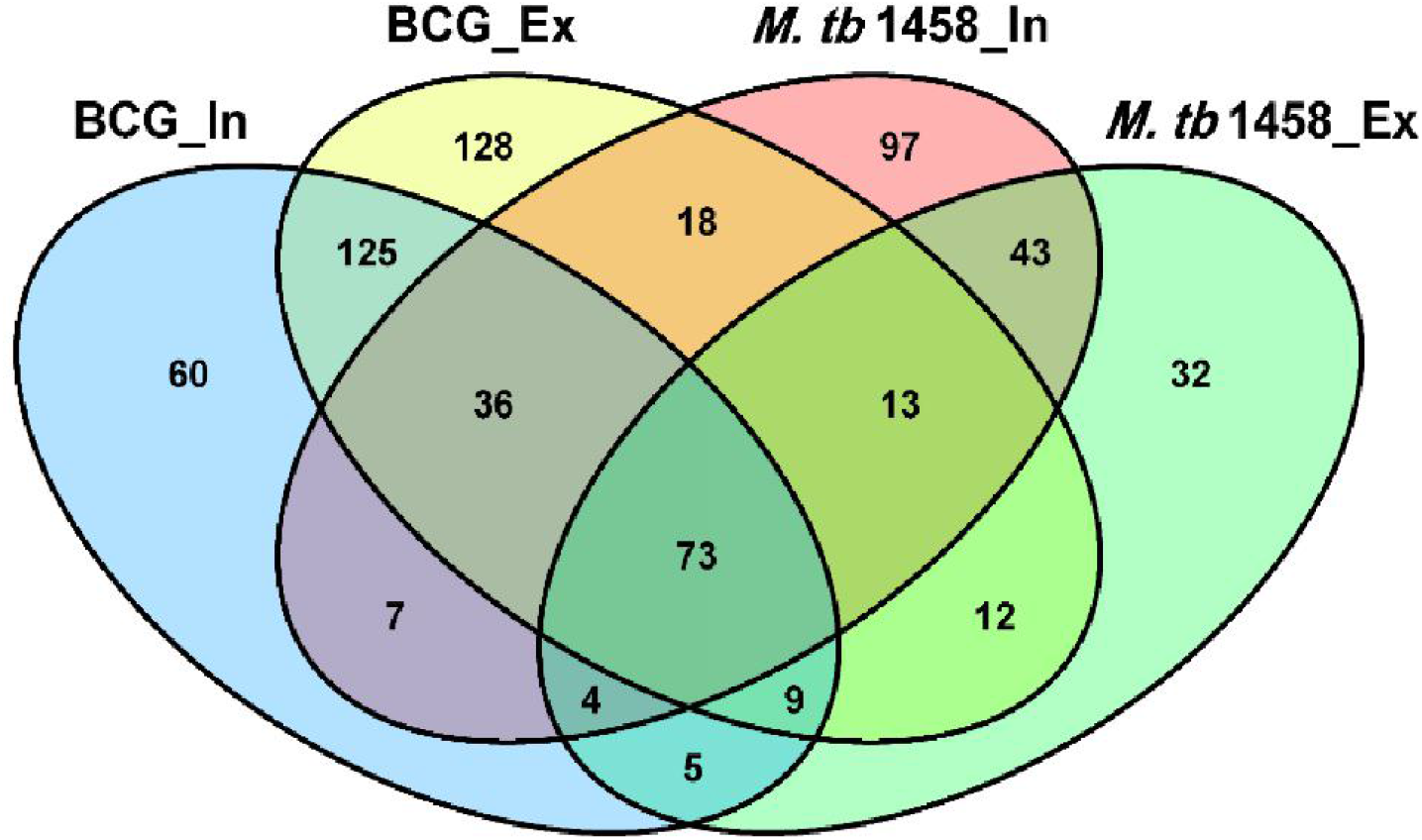
Venn diagram of the overlap after BCG and M. tb 1458 ncRNA sequences are compared. In stands for intracellular, Ex stands for extracellular.

### ncRNA annotation

Rfam and BRSD database were used to annotate ncRNAs. 96.73% in BCG and 96.28% in *M. tb*1458 were novel ncRNAs. For the annotated ones, besides very small part was rRNAs and tRNAs, others were reported sRNA with regulatory functions, like sRNA G2, C8, Glycine, mraW, 6C, ydaO-yuaA (Mandal, et al. 2004; Weinberg, et al. 2007; Kwon and Strobel 2008; Arnvig and Young 2009; Weinberg, et al. 2010; Nelson, et al. 2013; Mai, et al. 2019)(Fig.5). All the annotated ncRNAs were removed; remaining differential ncRNAs were used to predict the target and analyzed the pathway enrichment.

**Fig.5.**
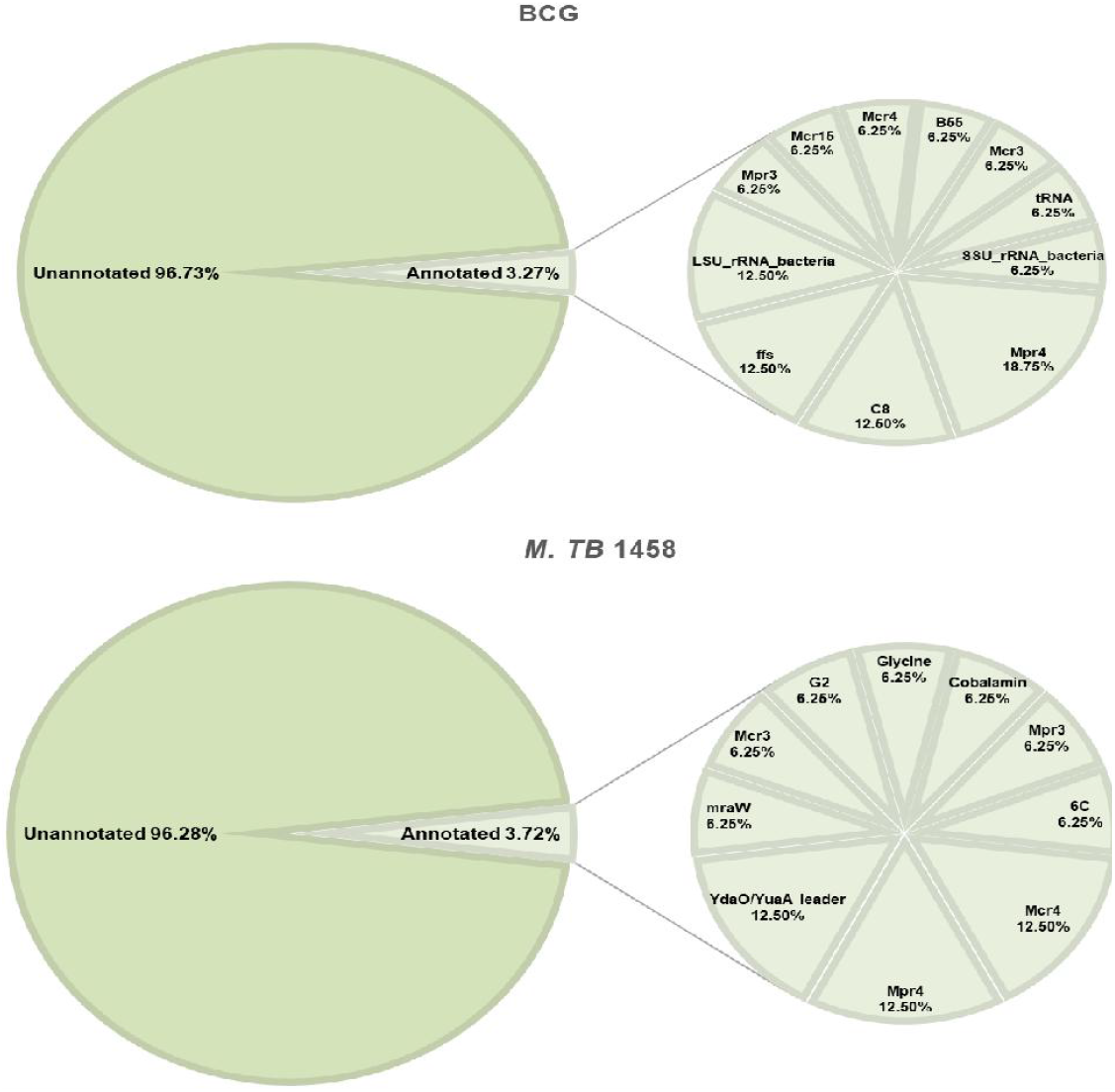
The annotation results of BCG and M. tb 1458 ncRNAs by Rfam and BRSD sRNA database.

### Target gene predication and enrichment analysis

IntaRNA, CopraRNA and TargetRNA2 were used to predict the target genes. Genes that predicated by all three tools were used to do the following analysis. Part of the ncRNA target genes were listed in table 1. In all listed 9 ncRNAs, target genes of 8 were related with lipid metabolism, such as lipid protein *lpqK*, *lpp and lprQ*, fatty acid metabolism related to the enzyme enyl coenzyme A hydrase *echA18* and *echA21*, Acetyl-coa dehydrogenase *fadE3*, *fadE16*, *fadD18*, *fadD28* and *fadE29*.

**Table 1.**
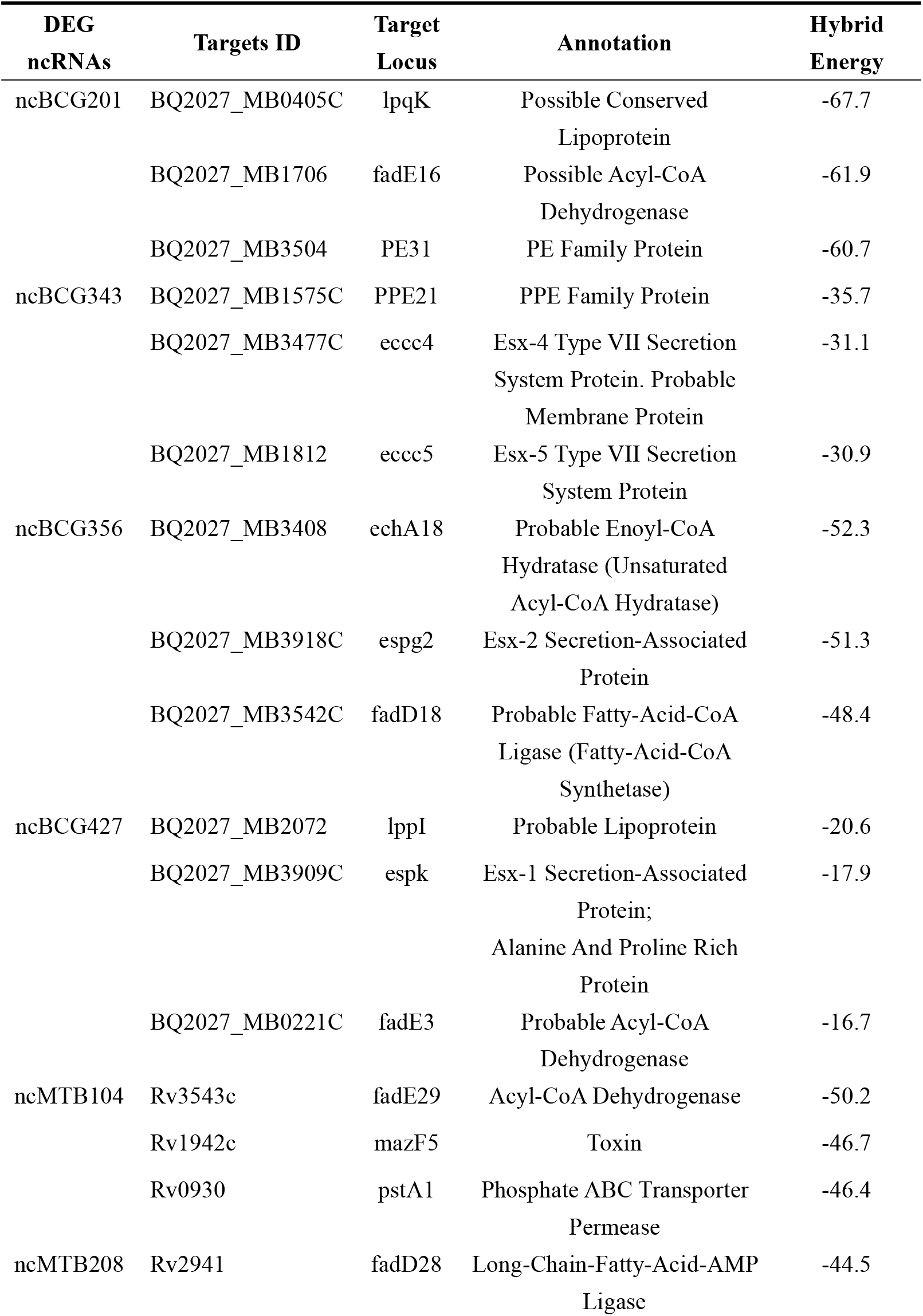
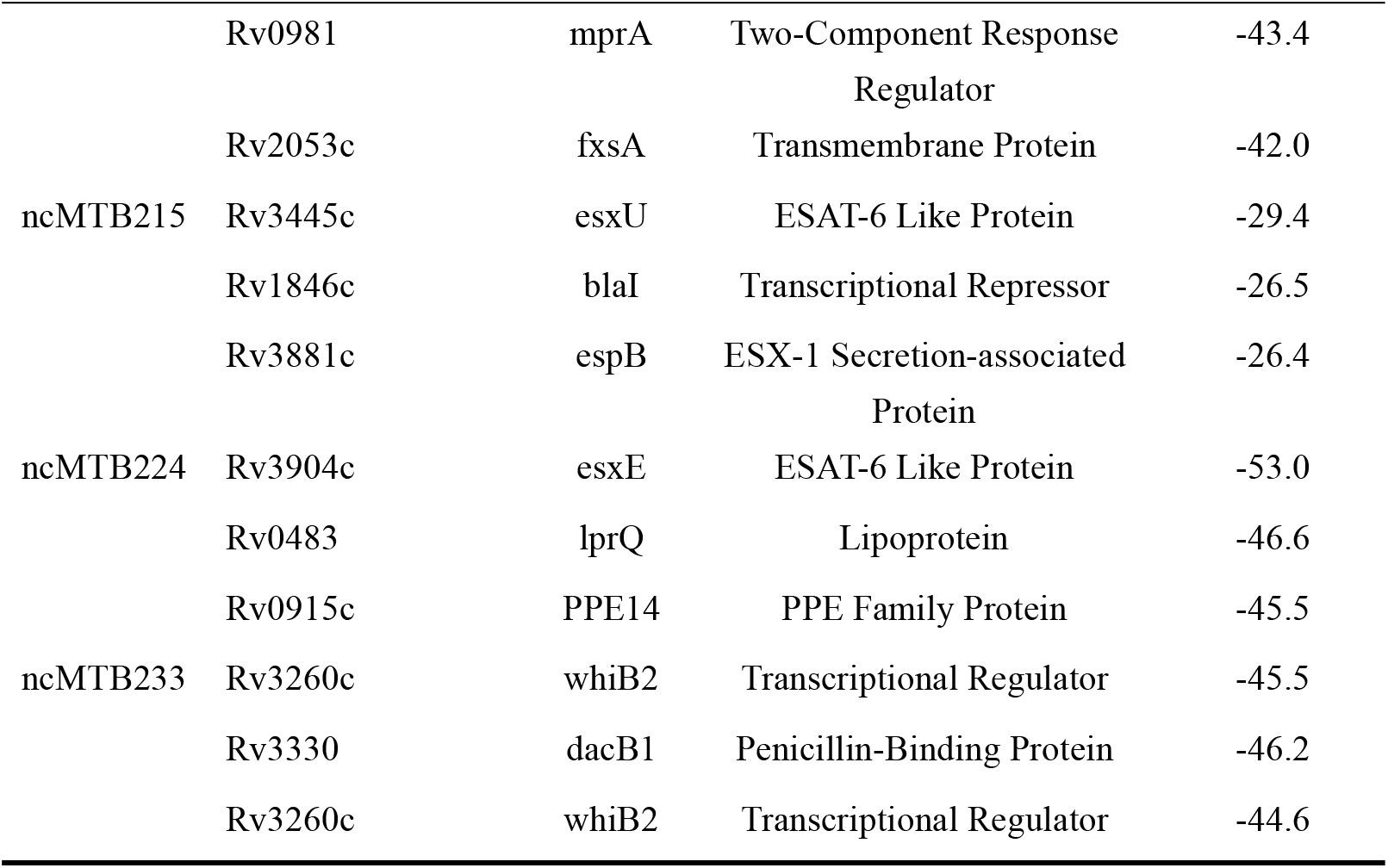
Important target genes of some DEG ncRNAs.

For the GO enrichment analysis, the most prevalent observed was biological process and molecular function, followed by cellular component (Fig. 6A). In Top 15 ncRNAs Terms in biological process, growth, pathogenesis, growth of symbiont in host, transmembrane transport, were all related with *mycobacterium* infection and intracellular survival. Among top 15 Terms in Molecular Function, fatty-acyl-CoA binding and acyl-CoA dehydrogenase activity were strongly related with *mycobacterium* fatty acid metabolism. ncRNAs in cellular component were significantly lower than other two groups.

**Fig.6.**
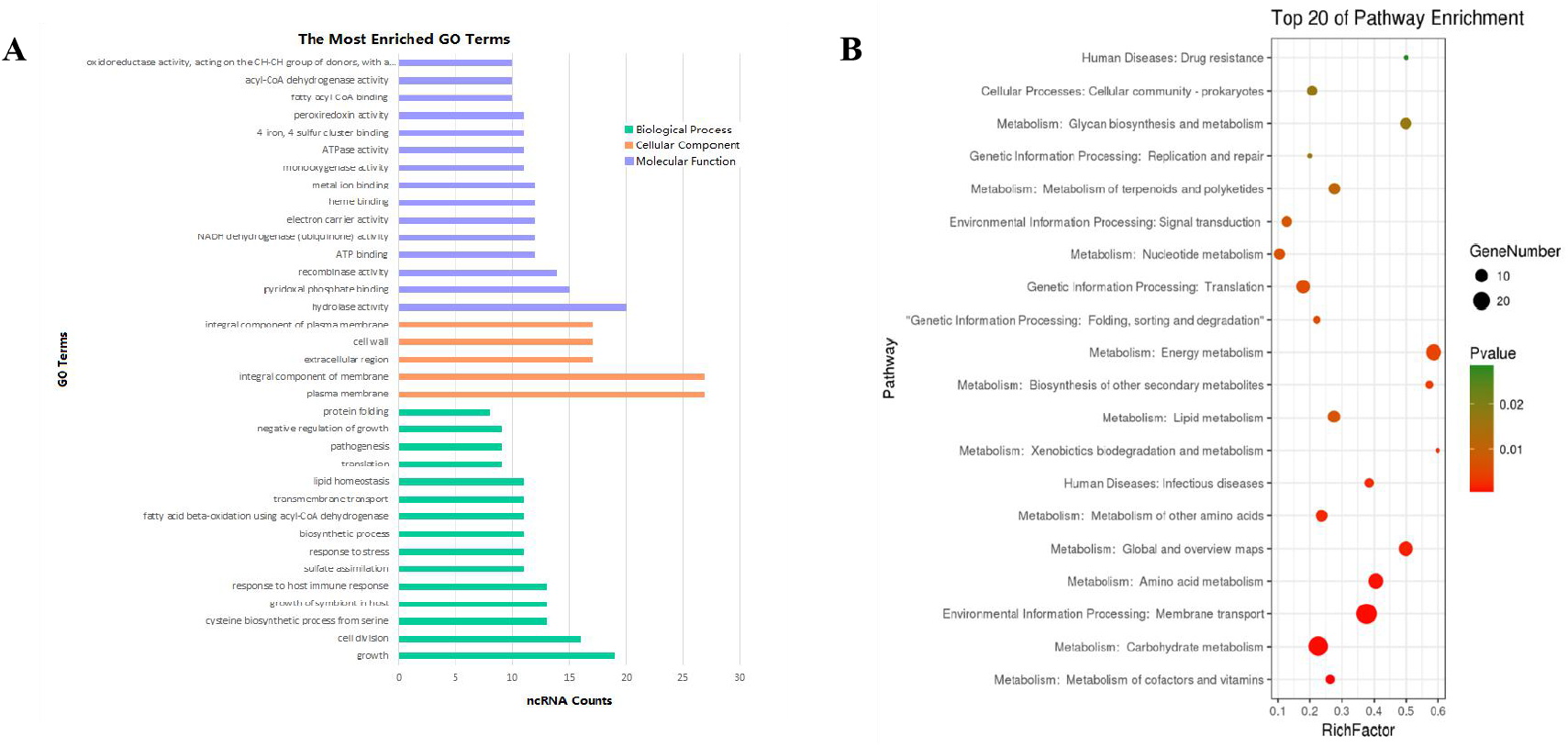
Enrichment analysis of differential expressed ncRNA (|FC|≥1.5). (A)The GO pathways (p≤0.1) of target that is most enriched in biological processes, cell component, and molecular functional. (B) Top 20 KEGG enrichment results (p≤0.1) of target genes.

To analyze the differential ncRNA target genes using the KEGG pathway, the top 20 were mainly engaged in bacterium metabolism, human diseases, genetic information processing and environmental information processing (Fig.6B). Interaction network of pathways were presented in supplemental figures. Because carbohydrate supply energy for bodies, carbohydrate metabolism network was no doubt the most complicated one, almost all the subgroup had ncRNAs, so we don’t focus on it. As lipid is one of the three major nutrients in life, and is essential for the bacteria survival, and the functional enzymes related to fatty acid metabolism have an important impact on the virulence regulation of *M. tuberculosis*, here we mainly focus on the lipid metabolism, both synthesis and degradation. ncBCG314 is related with glycerophospholipid metabolism, glycerolipid metabolism and fatty acid metabolism; ncBCG150 is related with glycerolipid metabolism, fatty acid metabolism and fatty acid degradation; ncBCG428 and ncBCG356 are associated with fatty acid metabolism, fatty acid degradation and synthesis and degradation of ketone bodies; ncBCG285 and ncBCG112 are related with glycerophospholipid metabolism, fatty acid metabolism and synthesis and degradation of ketone bodies; ncBCG98, ncBCG236, ncBCG337, ncMTB248, ncMTB233 were related with both glycerophospholipid and glycerolipid metabolism; ncMTB329 related with both synthesis and degradation of ketone bodies and glycerolipid metabolism (sFig.1).

For sRNAs that related with human diseases, ncBCG184, ncBCG285, ncBCG156, ncBCG341, ncBCG126, ncMTB64, ncMTB174, ncMTB165, ncMTB258 and ncMTB254 are engaged in tuberculosis. Among that, ncBCG285 and ncMTB165 also related to biosynthesis, which may indicated that those two sRNAs were participated in the drug resistant *mycobacterium* infection (sFig.2).

In genetic and environmental information processing pathways, number of ncRNAs in ribosomal pathway was the highest. Besides that, pathways about bacterium environmental processing including membrane transportation and two-component system were also included, membrane transportation pathway including ABC transporter and bacterial secretion system. Many ncRNAs were cross enriched to more than one pathways, like ncBCG429 was engaged in mismatch repair, microbial metabolism in diverse environments and two-component system; ncBCG34 was involved in microbial metabolism in diverse environments, two-component system and sulfur relay system; ncBCG403 related with two-component system, protein export and bacterial secretion system; ncBCG186 was related with sulfur relay system, protein export and bacterial secretion system; In *M. tb* 1458, more ncRNAs participate in multi pathways, like: ncMTB147 related with protein export, bacterial secretion system, nucleotide excision repair and quorum sensing; ncMTB258 involved in protein export, bacterial secretion system and quorum sensing; ncMTB206 involved in nucleotide excision repair, mismatch repair and sulfur relay system; ncMTB211, ncMTB20 and ncMTB164 were related to both microbial metabolism in diverse environments and ribosome (sFig.3).

### Identification of stress regulation related ncRNAs

Seventy-five differential expression ncRNAs (including 37 BCG ncRNAs and 38 *M. tb* 1458 ncRNAs) with potential biological functions predicated by biological information were selected and tested using relative real-time PCR in 6 stress culture models. 13 BCG ncRNAs and 14 *M. tb* 1458 ncRNAs were significantly expressed in stress environment. Differential expressed ncRNAs in different stress culture model were presented in sFig.4. Data showed that, ncBCG177, ncBCG181, ncBCG343, ncBCG356, ncMTB101, ncMTB126, ncMTB204 and ncMTB233 were differential expressed in all 6 stress environments. ncBCG368, ncBCG369, ncBCG378, ncBCG379, ncMTB104, ncMTB162, ncMTB208, ncMTB215, ncMTB221, ncMTB224 were differential expressed in 5 stress environments. ncBCG173, ncBCG201, ncMTB209, ncBCG349, ncBCG367, ncMTB214 and ncMTB239 were differential expressed in 4 stress environments. ncBCG390, ncBCG427, ncMTB127 were differential expressed in 3 stress environments. ncMTB228 was differential expressed in 2 stress environments.

For those ncRNAs, ncMTB162 was an antisense sRNA, ncBCG343, ncBCG378 and ncMTB208 were adjacent to the 5’UTR of downstream mRNA, others were located in the gene intergenic regions (sFig.5). Secondary structure prediction results showed that the loop of ncBCG177, ncBCG181, ncBCG201, ncBCG343, ncBCG356, ncBCG378, ncBCG427, ncMTB104, ncMTB162, ncMTB204, ncMTB208, ncMTB215, ncMTB224 and ncMTB233 were between 4 to 14bp, and had high GC content, indicated that those sRNAs can bind to target mRNA more precisely and firmly(sFig.6).

### Phenotype testing of recombinant *M. smegmatis*

Fourteen sRNAs (including 7 ncBCG RNAs and 7 ncMTB RNAs) with less than 100bp were overexpressed successfully in *M. smegmatis* (Fig.7). Recombinant MS Vector was used as control. For the growth curves, MS_ncBCG201 showed significantly greater growth speed since 24h (p<0.05), especially significantly higher from 94h (p<0.0001); MS_ncBCG427 showed a significantly increase in growth since 84h (p<0.0001), but quickly entered the plateau phase at 120h, then grow slowly since 144h (p<0.0001); MS_ncBCG378 and MS_ncBCG 356 also showed a higher growth speed since 60h and 84h, respectively (p<0.001) (Fig.8A). For recombinant *M. smegmatis* of ncMTB sRNAs, MS_ncMTB162 grows significantly quicker from 24 h, respectively (p<0.0001), MS_ncMTB 233 and MS_ncMTB224 showed a similar trend, grows slowly since 48 h (p<0.001, p<0.01), MS_ncMTB215 grows significantly quickly since 84h and lasted for a long time (p<0.0001) (Fig.8B).

**Fig.7.**
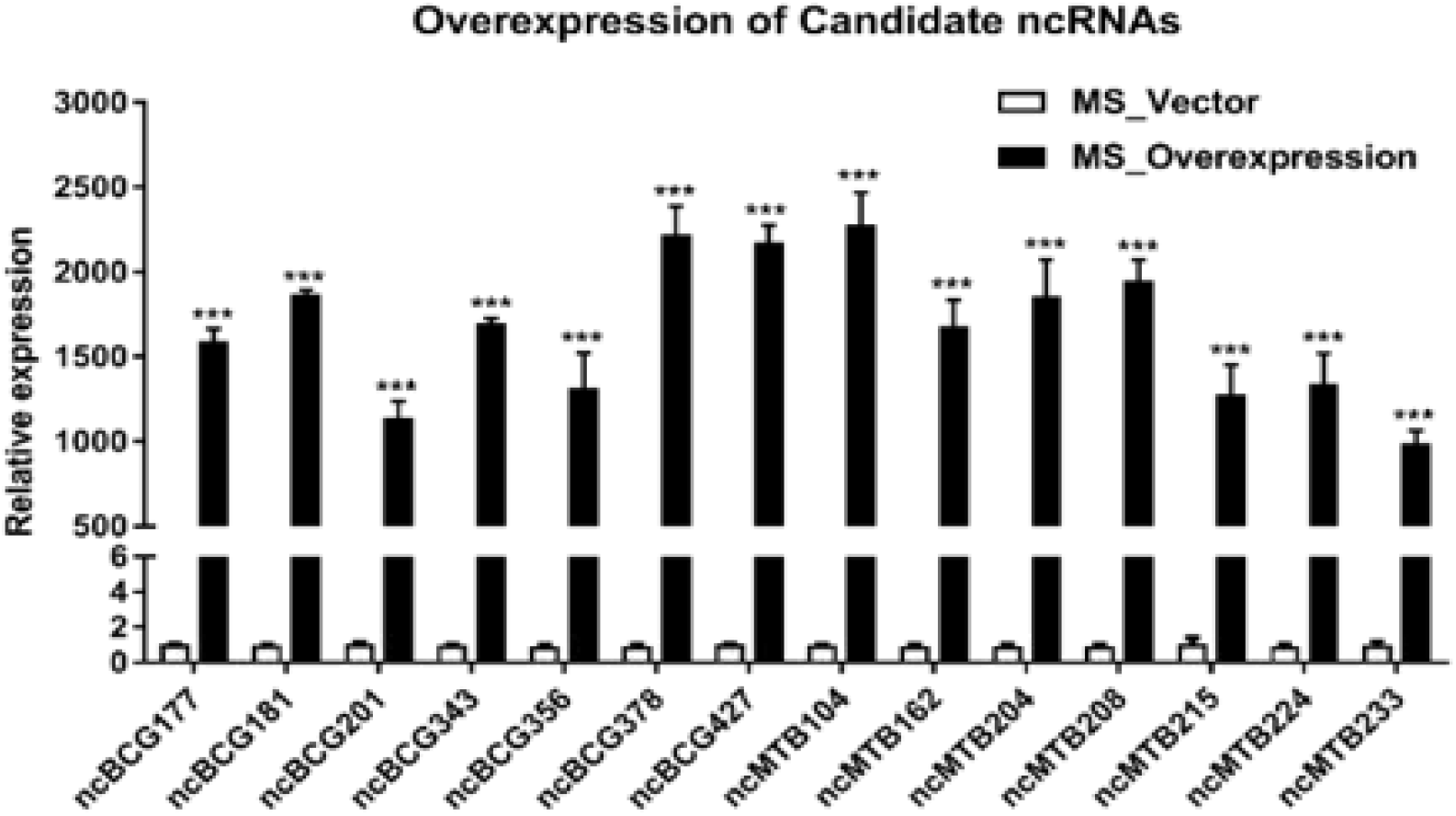
Expression of candidate sRNAs in each overexpressing M.S strain. The data shown are the mean ± SD of three replicate samples. t-test was used for statistical analysis, * represents p<0.05, ** represents p<0.01, and *** represents p<0.001.

**Fig.8.**
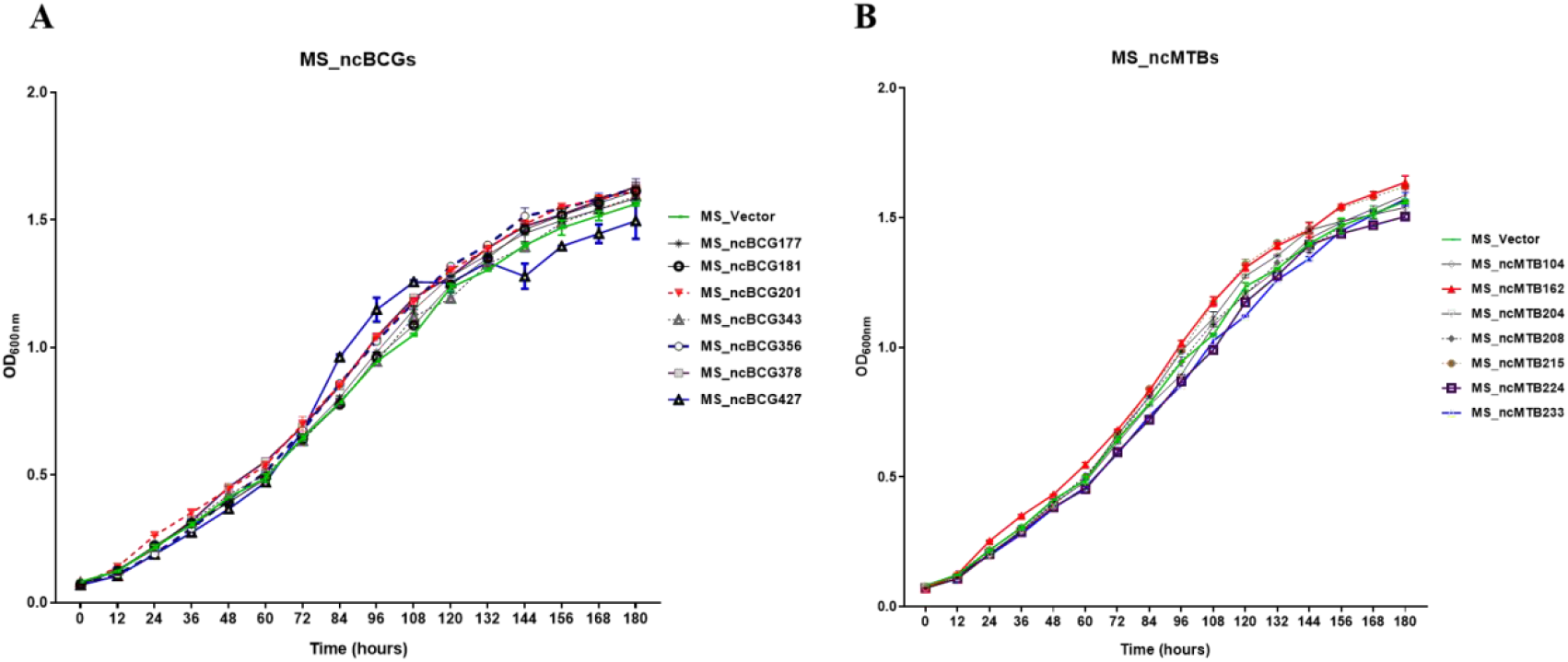
Growth curve of candidate sRNA overexpressing *mycobacterium smegmatis* MC^2^ 155 strains.(A)ncBCG sRNA overexpressing *mycobacterium smegmatis* MC^2^ 155 strains; (B) ncMTB sRNA overexpressing *mycobacterium smegmatis* MC^2^ 155 strains.

Single colony morphology was also tested. Recombinant MS_Vector showed a smooth and moist morphology, MS_ncBCG356, MS_ncBCG177, MS_ncBCG181, MS_ncBCG201, MS_ncBCG378, MS_ncBCG343 and MS_ncMTB224 showed a thinner colony. Except MS_ncMTB224, other 6 strains had significantly larger colony than control (Fig.9). The most different one is MS_ncBCG427, which showed a significantly smaller and rough colony (Fig.10A and B); followed by MS_ncBCG215, which is larger but also rough (Fig.10A and C). Those two strains had very obviously folds whose ridge is obviously higher than control, which indicated that sRNA ncBCG427and ncMTB215 may affect the bacterial growth by regulate the cell wall components.

**Fig.9.**
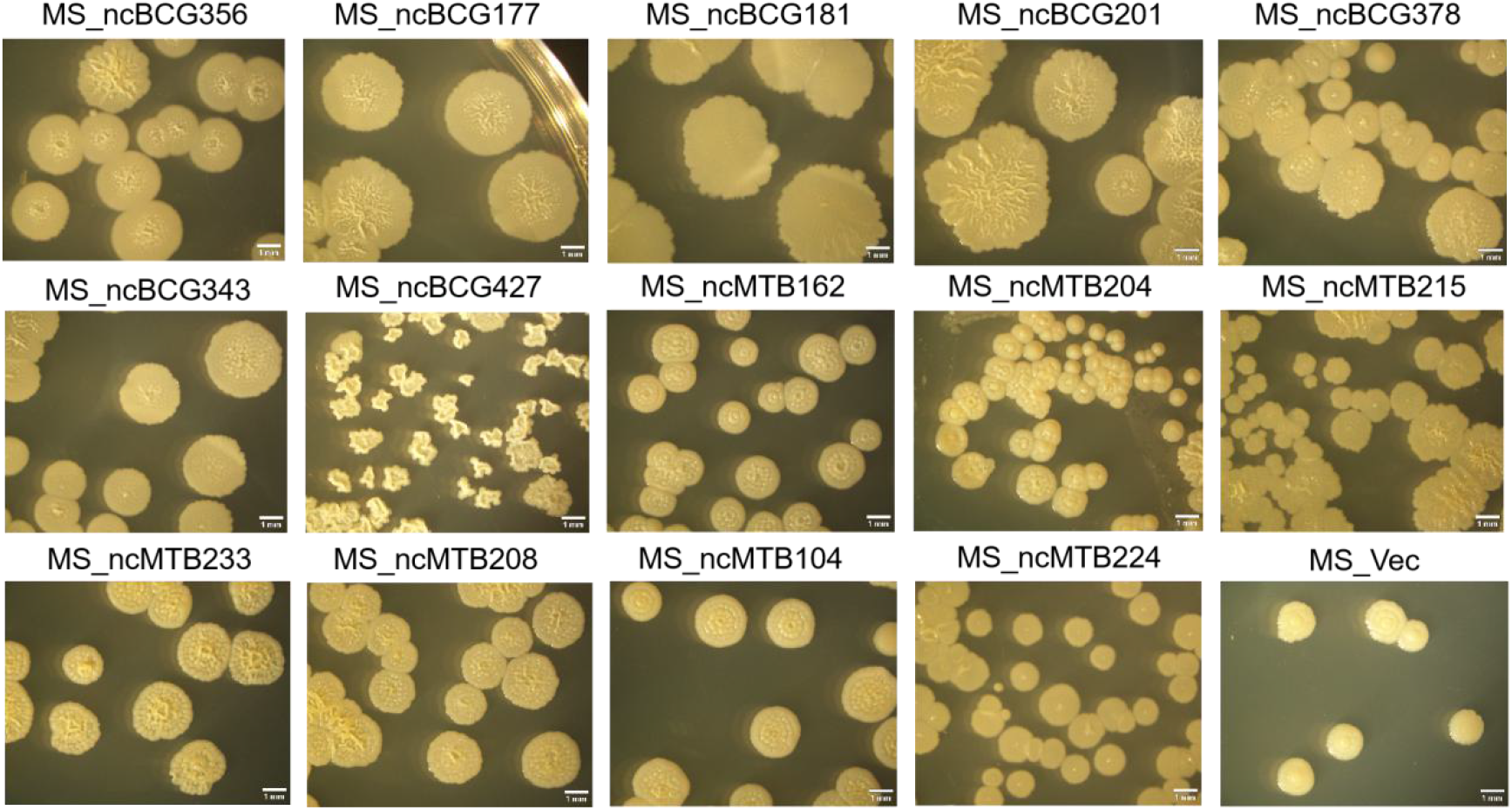
single colony morphology of candidate sRNA overexpressing *mycobacterium smegmatis* MC^2^ 155 strain.

**Fig.10.**
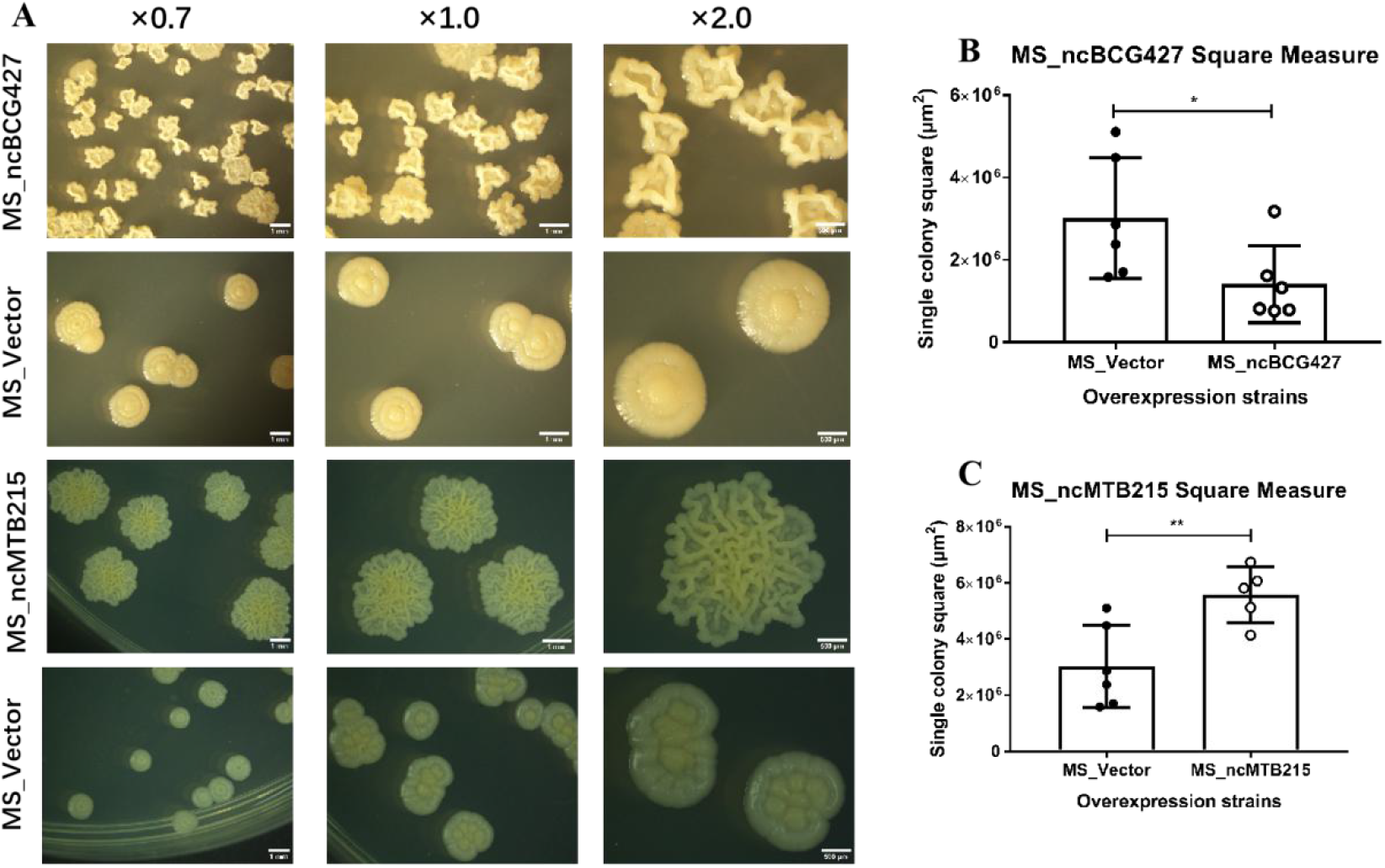
Single colony morphology and square measure of overexpressing *mycobacterium smegmatis* MC^2^ 155 strains MS_ncBCG427 and MS_ncMTB215. (A) Single colony morphology of different strains; (B) square of MS_ncBCG427; (B) square of MS_ncMTB215.The two-way ANOVA method was used for statistical analysis, where * represented p<0.05, ** represented p<0.01, and *** represented p<0.001.

### MIC of recombinant *M. smegmatis*xs

MIC of RFP, INH, PAS, SM, EMB and LVX were tested. Data showed that MS_ncBCG427 decreased the sensitivity to INH and PAS, but increase the sensitivity to RFP and LVX; MS_ncBCG181, and MS_ncBCG201 decrease the sensitivity to PAS but increased the sensitivity to RFP; MS_ncBCG378 and MS_ncBCG343 increased the sensitivity to both RFP and SM; MS_ncBCG177, MS_ncMTB215, MS_ncMTB233, MS_ncMTB104 and MS_ncMTB224 showed a higher sensitivity to RFP, and MS_ncBCG356 was less sensitivity to PAS (Table 2).

**Table 2.** Drug sensitivity of recombinant *M. smegmatis* strains overexpressing sRNA of BCG/*M. tb* 1458.

| 1458 |  |  |  |  |  |
| --- | --- | --- | --- | --- | --- |
| MIC (μg/mL) | MS_Vector | MS_ncBCG356 | MS_ncBCG177 | MS_ncBCG181 | MS_ncBCG201 |
| INH | 4 | 4 | 4 | 4 | 4 |
| RFP | 32 | 32 | 16 | 16 | 16 |
| SM | 0.5 | 0.5 | 0.5 | 0.5 | 0.5 |
| EMB | 64 | 64 | 64 | 64 | 64 |
| LVX | 0.125 | 0.125 | 0.125 | 0.125 | 0.125 |
| PAS | 512 | 1024 | 512 | 1024 | 1024 |
| MIC (μg/mL) | MS_ncBCG378 | MS_ncBCG343 | MS_ncBCG427 | MS_ncMTB162 | MS_ncMTB204 |
| INH | 4 | 4 | 8 | 4 | 4 |
| **RFP** | 8 | 16 | 16 | 32 | 32 |
| **SM** | 0.25 | 0.25 | 0.5 | 0.5 | 0.5 |
| **EMB** | 64 | 64 | 64 | 64 | 64 |
| **LVX** | 0.125 | 0.125 | 0.0625 | 0.125 | 0.125 |
| **PAS** | 512 | 512 | 1024 | 512 | 512 |

| **MIC (µg/mL)** | **MS_ncMTB215** | **MS_ncMTB233** | **MS_ncMTB208** | **MS_ncMTB104** | **MS_ncMTB224** |
| --- | --- | --- | --- | --- | --- |
| **INH** | 4 | 4 | 4 | 4 | 4 |
| **RFP** | 16 | 16 | 32 | 16 | 15 |
| **SM** | 0.5 | 0.5 | 0.5 | 0.5 | 0.5 |
| **EMB** | 64 | 64 | 64 | 64 | 64 |
| **LVX** | 0.125 | 0.125 | 0.125 | 0.125 | 0.125 |
| **PAS** | 512 | 512 | 512 | 512 | 512 |

### Biofilm formation ability of recombinant *M. smegmatis*

The biofilm differences between sRNA overexpressed strain and MS_Vector was compared. Data showed that except MS_ncBCG177, MS_ncBCG201 and MS_ncMTB204, other recombinant strains all had significantly different biofilm forming ability compared with MS_Vector. The biofilms of MS_ncBCG356, MS_ncBCG181, MS_ncBCG378, MS_ncBCG343, MS_ncMTB215 and MS_ncMTB104 were sparse and easy to be broken when under shaking; although MS_ncMTB162 and MS_ncMTB233 had lower biofilm forming ability, it was still robust and not fragile. MS_ncBCG427, MS_ncMTB208 and MS_ncMTB224 had higher biofilm forming ability compared with MS_Vector (Fig.11).

**Fig.11.**
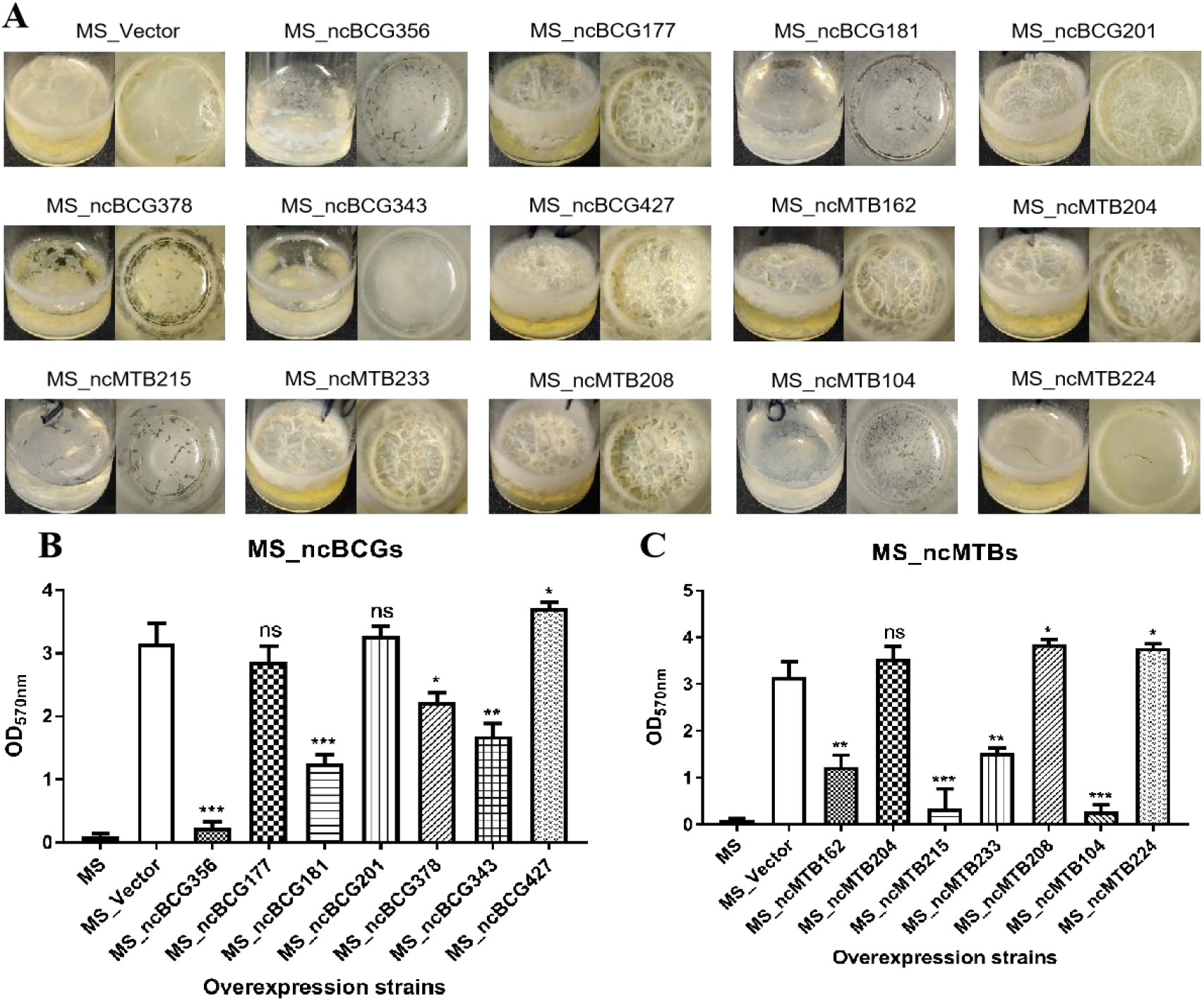
Biofilm formation ability of the candidate sRNA over-expressing strains. (A) Biofilm morphology; (B) Crystal violet staining results of MS_ncBCGs; (C) Crystal violet staining results of MS_ncMTBs; compared with MS_vector. t-test was used for statistical analysis, * represents p<0.05, ** represents p<0.01, and *** represents p<0.001.

### Invasion of A549 ability of recombinant *M. smegmatis*

As MS_ncBCG427 had totally different colony and biofilm morphology, changed the resistance to 4 drugs, and grow significantly quickly from the log phase, the invasion ability was then tested. Result showed that MS_ncBCG427 had lower invasion ability compared with MS_control strain (Fig. 12).

**Fig.12.**
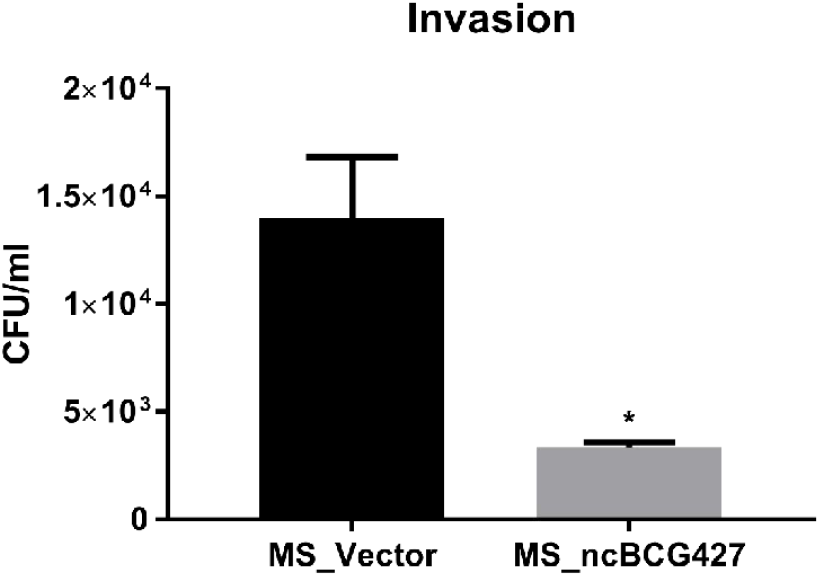
The invasion ability of overexpressing *mycobacterium smegmatis* MC^2^ 155 strain MS_ncBCG427. The t-test was used for statistical analysis, where * represented p<0.05, ** represented p<0.01, and *** represented p<0.001.

### Target gene prediction and identification in *M. smegmatis*

163 target genes were enriched to cytoplasm pathway, 133 in plasma membrane pathway, 51 in ATPase activity pathway, 43 were enriched to acyl-CoA dehydrogenase activity and translation, respectively, 41 in structural constituent of ribosome pathway, 32 engaged in ribosome pathway, 27 in amino sugar and nucleotide sugar metabolism pathway, 10 in fluorobenzoate degradation pathway, 9 in alanine racemase activity pathway (Table S3). Consider the colony and biofilm morphology and the predict target genes GO enrichment, we inferred that ncBCG427 may be involved in the regulation of acyl CoA dehydrogenase activity and plasma membrane formation; Real-time PCR (RT-PCR) was used to identify the target gene expressions. Four genes (MSMEG_0578, MSMEG_4076, MSMEG_4096 and MSMEG_4844) in acyl-CoA dehydrogenase activity pathway and 6 genes (MSMEG_0083, MSMEG_1875, MSMEG_2461, MSMEG_3547, MSMEG_3601 and MSMEG_5318) in plasma membrane pathway were upregulated, indicated that ncBCG427 may have a negative regulatory effect on its target mRNA (Fig.13).

**Fig.13.**
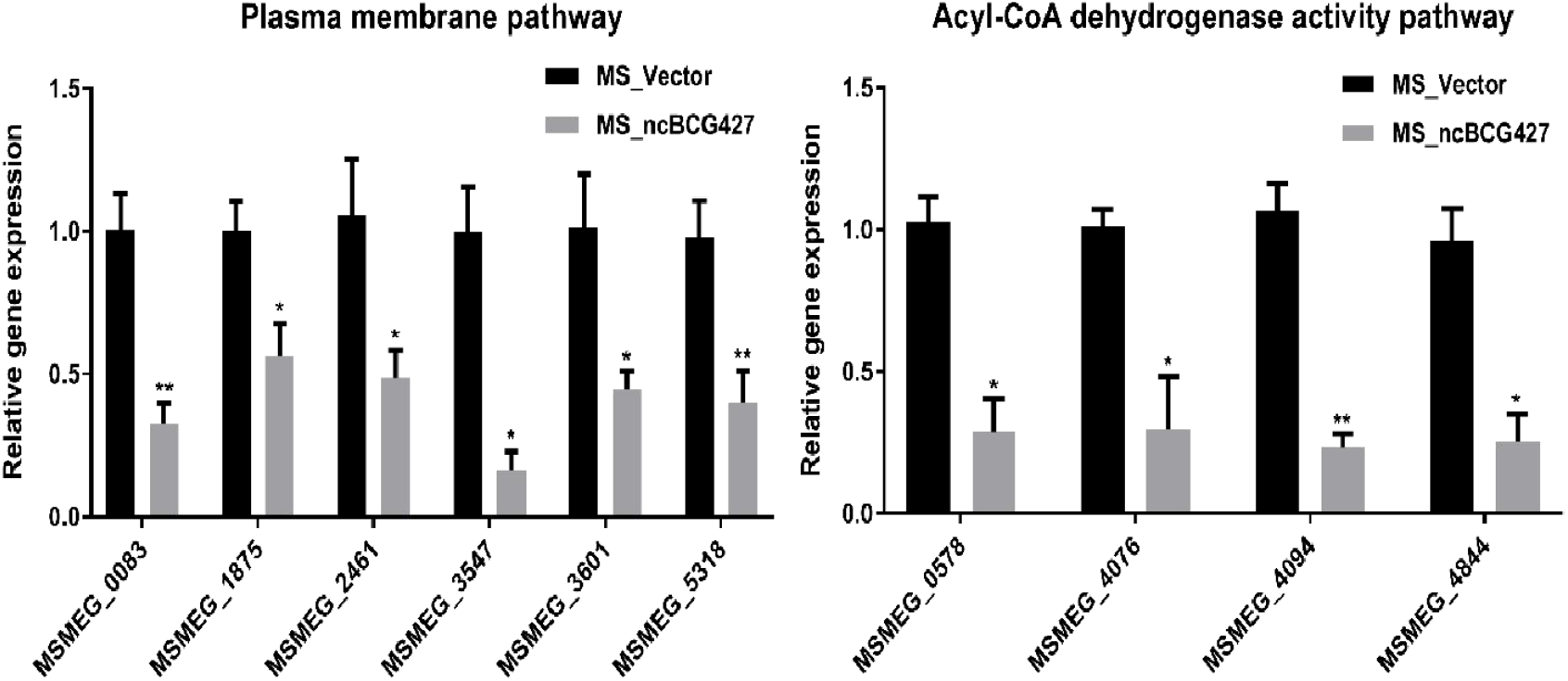
The differential expression of MS_ncBCG427 predicted target genes in M. smegmatis Mc2 155. The t-test was used for statistical analysis, where * represented p<0.05, ** represented p<0.01, and *** represented p<0.001.

## Discussion

The binding of sRNA to its target not only prevents ribosome binding but also initiates the cleavage of RNA-RNA interaction. In Staphylococcus aureus, for example, sRNA controls expression of several pathogenesis-related genes binding to the target RNA (Ostrik, et al. 2021). Initial analysis of the *M. tb* transcriptome by RNA-seq reveal the presence of at least 15 sRNAs (Westermann 2018). Currently, many more regulatory sRNA have been identified in *Mycobacterium* using saturated transposon mutagenesis (Budell, et al. 2020). More recent studies have evidenced the importance of sRNA in pathogenesis. Therefore, sRNA represents a regulatory mechanism to provide an adaptive response to fast changing environments, and potentially are quite important in slow growth organisms.

In the current research, we attempted to comparatively identify regulatory mechanisms in the human pathogen M. tb and the animal pathogen *M.bovis*. By exposing both microorganisms to conditions encountered in the host, we identify many new sRNA sequences.

### A RNA-seq comparison between extracellular and intracellular of BCG and *M. tb*

We used BCG and *M. tb* 1458 strains to infect THP-1, in order to characterize the ncRNA differences both inside and outside of infected cells. Past study has demonstrated that *M. tb* 1458 was capable to infect macrophages more efficiently, and have an enhanced virulent compared to *M. bovis* BCG (Xiong, et al. 2017; Zhang, et al. 2018). The ncRNAs obtained from the extracellular BCG was more abundant than the one obtained from the intracellular environment. In comparison, there was less ncRNAs in extracellular than intracellular *M. tb*. The observation may be related to the fact that virulent *M. tb* 1458 has higher macrophage uptake. For all the annotated ncRNAs, around 20% were rRNA and tRNA, which are essential for ribosome and translation (Ravishankar, et al. 2016), and generally are not involved in post transcriptional level of regulation. Mcr3 (Mpr7) and Mpr4 were expressed in both BCG and *M. tb 1458* in our study. Mcr3, which is also reported as a conserved sRNA in *M. smegmatis*, may be involved in the regulation of functions in different mycobacteria (DiChiara, et al. 2010). C8, in fact 4.5S RNA, is an antisense sRNA which is known to be trimmed of the first 90 nucleotides and it is highly conserved in mycobacteria as well as in more distantly related bacteria, such as Rhodococcus, Corynebacteria and Nocardia (Arnvig and Young 2009). B55 and G2 are two sRNAs reported conserved in *Mycobacterium tuberculosis* complex. B55 may be part of the 3′UTR of the Rv0609A mRNA rather than being an sRNA, G2 is a trans‐ encoded sRNA with more than one 5’end, and may act as a possible SigC promoter upstream of one of the 5’ends, which can downregulate the growth of *mycobacterium* (Arnvig and Young 2009). Glycine riboswitch was only reported in *Bacillus subtilis*, regarded as a unique cis-acting mRNA element that contains two tandem homologous glycine-binding domains, and act on a single expression platform to regulate gene expression in response to glycine (Babina, et al. 2017). In fact, riboswitching-binding metabolites have been successfully used to inhibit growth of non-pathogenic *Bacillus subtilis* in vitro (Ritchey, et al. 2020). 6C is a conserved sRNA in high GC Gram-positive bacteria including *Mycobacterium*; it has many cellular functions via binding to multiple mRNA targets through the C-rich loops in *Mycobacterium*, and regulates DNA replication and protein secretion (Mai, et al. 2019). YadO is a riboswitch, which regulate the resistance to enzyme-mediated degradation (Meehan, et al. 2016).

Our results showed that many sRNAs expressed intracellular at the timepoint examined, have functions that are not completed understood, and future studies will have to address the gap in knowledge.

It is also important to consider that cell-to-cell heterogenicity in RNA expression may occur.

### Biofilm formation ability

Biofilm plays an important role in *mycobacterium* pathogenicity, and also drug resistance. Biofilm formation ablility influence the bacterium existence in adverse environments. In mycobacterial, many cell surface molecules especially many lipids were associated with biofilm formation, such as glycopeptidolipids, poly-a-L-glutamine, mycolic acid, PPE family proteins (Xiang, et al. 2014; Crotta Asis, et al. 2021). *M.tuberculosis* and *M.bovis* form cords, and biofilm inside the granolomatous lesion as well intracellularly. Role of the biofilm in those conditions is not well understood and may relate to the absence of nutrients, although inside cells the synthesis of trehalose-6-6-dimycolate (TDM or cord factor) maybe reduced due the shift in metabolism related to the abundance of glucose, leading to the systhesis glucose monoglycolate, instead of TDM (Hunter, et al. 2006).

Here the differential expressed sRNAs ncBCG201 (RPKM fold of extracellular/intracellular was 2.24), ncBCG343 (unique expressed intracellularly) and ncMTB224 (unique expressed in extracellular *M. tb* 1458) were predicted target to PE/PPE family genes. Past study reported PE/PPE genes are mainly found in virulent mycobacteria (Li, et al. 2019), that agree with our differential expressing results; as PE/PPE proteins are important for the *mycobacterium* growth, and could be differentially expressed under a variety of conditions (Li, et al. 2019), which suggested that ncBCG201, ncBCG343 and ncMTB224 might participate in the pathogenesis and growing bacterium.

Recombinate strain of those three sRNAs all showed a signifncantly change in growing rate (not obviously for ncBCG343), small colonies, different biofilm forming abilities, which were all associalted with the virulence of bacterium (Ramsugit and Pillay 2014; Cardona 2018), confirmed that ncBCG201, ncBCG343 and ncMTB224 have potential regulating pathogenesis ability. In addition, all three RNAs changed the drug resistant ability, indicated that they may also target in some genes that associated with the drug resistance.

### Lipid metabolism regulation ability

Lipids constitute a large percent of the mycobacteria outstructure. Many of the virulent Mycobacteria, such Mycobacteria avium subsp paratuberculosis, change the lipids of the cell wall following infection (Alonso-Hearn, et al. 2010). The current results show that lipid metabolism is strongly associated with many target genes of differential expressed sRNAs, such as *lpqK*, *lppI,lprQ*, Enoyl-CoA Hydratase *echA18* and *21*, Acyl-CoA Dehydrogenase fadE3, 16, fadD18, 28 and 29.

The composition and organization of the mycobacterial cell envelope is very complex, which is a distinctive feature of the *Mycobacterium* genus. The cell envelop plays multiple roles in *mycobacterium* infection, including the modulation of the phagosome-maturation, granuloma biogenesis and is involved in the ability of the bacteria to adapt to all kinds of environmental stresses (Gago, et al. 2018). Lipid metabolism, especially fatty acids metabolism, is essential in the *mycobacterium* life cycle, and can affect bacterial virulence and drug tolerance during infection (Nazarova, et al. 2017).

*M. tb* encodes five type VII secretion systems (ESX-1-5), whose genes are arranged in highly conserved clusters. Here, espk (ncBCG427), esxU and espB (ncMTB215), esxE (ncMTB224), associated with ESX-1, can modulate necrosis, NOD2 signaling, type I interferon production, and autophagy (Wong 2017). Eccc5 (ncBCG343), an Esx-5 Type VII secretion system protein, which is essential for growth and participate on the cell surface and cell envelope fractions transportation (Ates, et al. 2015); espg2 is a Esx-2 secretion-associated protein, although its function is unclear.

As target genes of differential expressed ncRNA were mainly focus on participate the biological progress not be the cell component, although they are multiple functions, GO enrichment analysis still discover the top function was associated with lipid or membrane. This information also confirmed by our KEGG analysis.

### Ability to Adapt to the Environment

Tuberculosis is an old disease. During thousands of years, pathogens evolved many ways to survive within the host and to adapt the ever-changing environment, including extracellular and intracellular. In different nutritionally-defined environments, like lack of carbon and nitrogen sources, low tension of oxygen environment, lipid composition of the envelope will be changed, resulting in the alteration of growth rate, metabolic activity and resistance to antibiotics (Gago, et al. 2018). Under different stresses and environments, almost all sRNA recombinant expressed in *M. smegmatis* had some sort of change in bacterial adaptation. More specifically, ncBCG177, ncBCG 181, ncBCG 343, ncBCG 356, ncMTB204 and ncMTB233, not only changed the expression level in all of the six stress models, but altered the bacterial growth rate and colony morphology, suggesting that the expressed sRNAs encoded the ability to the bacterium to adapt to adverse environments. Among the strains, the *M.smegmatis* expressing ncBCG427 had the most different morphology among all strains. Its expression had effect of biofilm robustness, resistance to antibiotics and improved the ability of immune escape, changing pathogen virulence (Rastogi, et al. 2017; Wang, et al. 2019). Our observation also discover that ncBCG427 had change in susceptibility to 4 first-line drugs for tuberculosis INH, PAS, RFP and LVX, as well as decreased ability to invade host cells, compared to controls.

## Conclusion

Taken together, 490 ncRNAs in BCG and 349 ncRNAs in *M. tb1458* were identified by RNA-seq. In a subset of differentially expressed sRNAs that are infected in multiple stress conditions, many of them were associated with lipid metabolism. Among them, ncBCG427, was significantly down-regulated when BCG entered into macrophages, and was associated with increase of biofilm formation and changed in drug susceptibility. Then, reduction of virulence possibility depends on regulating lipid metabolism.ncBCG427, significantly down-regulated when *M. bovis* BCG is ingested by macrophages, is associated with biofilm formation in *mycobacterium*, results in changes in the pattern of drug susceptibility, and reduces virulence-related phenotype that depends upon lipid metabolism.

Many studies in the past have indicated the role of lipids in *M. tb* virulence, and support the observations that lipids are released from the bacterium in a constant manner, very likely indicating their crucial role in the survival of the pathogen. The understanding of the regulation of the lipid synthesis and release will provide important information about future drug targets.

## Acknowledgements

This research was supported by National Key Research and Development Program of China (2017YFD0500300), China Agriculture Research System of MOF and MARA, and National Distinguished Scholars in Agricultural Research and Technical Innovative Team.

